# Fructose appetition in “taste-blind” P2X2/P2X3 double knockout mice

**DOI:** 10.1101/2021.07.22.453428

**Authors:** Anthony Sclafani, Karen Ackroff

**Affiliations:** Department of Psychology, Brooklyn College of the City University of New York, Brooklyn, NY 11210, USA

**Keywords:** Sweet taste, glucose, saccharin, sucralose, flavor conditioning, postoral sugar reinforcement

## Abstract

Inbred mouse strains differ in their postoral appetite stimulating response (appetition) to glucose and fructose. For example, C57BL/6J (B6) and FVB strains learn strong preferences for a flavor (CS+, e.g., cherry) paired with intragastric (IG) glucose infusions, but only FVB mice learned to prefer a CS+ paired with IG fructose infusions. Consistent with these findings, “tasteless” B6 knockout (KO) mice missing the taste signaling protein TRPM5 learn strong preferences for a CS+ added to glucose solution as well as for unflavored glucose but weak or no preferences for a fructose-paired CS+ or unflavored fructose. The present experiment reports that “tasteless” P2X2/P2X3 double-knockout (P2X2/3 DKO) mice, unlike TRPM5 KO mice, learned strong preferences for a CS+ mixed with fructose as well as for unflavored fructose. Whether differences in genetic backgrounds or other factors account for the fructose appetition displayed by P2X2/3 DKO mice but not TRPM5 KO mice remains to be determined.

## 1. Introduction

Sugar preference in rodents is initially determined by sweet taste signaling but can be substantially modified by the nutrient’s postoral actions [4]. This is demonstrated by the ability of intragastric (IG) sugar infusions to enhance preferences for already preferred sweet solutions (e.g., cherry saccharin) or create preferences for initially avoided solutions (e.g., bitter sucrose octaccetate) [10;17]. Also, the initial preference for a nonnutritive sweetener (e.g., sucralose + saccharin mixture, S+S) over a sugar (e.g., glucose) can be reversed after the animal experiences the sugar’s postoral actions [21]. The postoral appetite stimulating actions of sugar, referred to as appetition [11], vary as a function of the type of sugar. In particular, glucose and glucose-containing carbohydrates (sucrose, maltose, maltodextrin) have stronger appetition effects than does fructose in rats and several inbred mouse strains [3;12;15;16]. For example, C57BL/6 (B6) mice trained to drink a flavored solution (CS+) paired with IG sugar and a different flavor (CS−) paired with IG water acquire a robust preference for a glucose-paired CS+ but no preference for a fructose-paired CS+ [12;24]. Yet, B6 mice learn to prefer a flavor added to an orally consumed fructose solution, which is attributed to a learned association between the CS+ flavor and the sweet taste of fructose, i.e., flavor-taste learning [14]. Unlike B6 mice, some mouse strains acquire preferences for a CS+ flavor paired with IG fructose (FVB mice) and learn to prefer a fructose solution to a S+S mixture (FVB, SWR mice) [8;20]. Nevertheless, these strains, like B6, acquire strong preferences for glucose over fructose, indicating that glucose has more potent appetition effects than fructose [8;19;20].

Further evidence for the differential postoral actions of glucose and fructose is provided by the sugar preferences displayed by sweet “tasteless” knockout (KO) mice missing the T1R3 sweet receptor component (T1R3 KO mice) or the TRPM5 sodium channel (TRPM5 KO mice). In 24-h sugar vs. water tests these KO mice were indifferent to dilute sugar solutions, but developed significant preferences for concentrated (8-32%) glucose but not fructose solutions [22;26]. The preference for the glucose solutions can be attributed to a postoral conditioned preference for the residual flavor properties (odor, texture) of the sugar solution. In contrast to these findings, a recent study reported that “tasteless” P2X2/P2X3 double knockout (P2X2/3 DKO) mice, which are missing ATP receptors required for taste cell signaling to gustatory nerves, displayed preferences for concentrated (15%) fructose as well as glucose and sucrose in 24-h sugar vs. water tests [2]. Here we provide additional evidence that fructose has postoral appetition actions in P2X2/3 DKO mice. In 2015 we reported that TRPM5 KO mice trained 24 h/day to drink a CS+ flavored (e.g., grape) fructose solution and CS− flavored (e.g., cherry) water displayed only a weak preference for the fructose-paired CS+ and no preference for unflavored fructose, which contrasts with the strong preferences they displayed for glucose and a glucose paired CS+ flavor [14]. Shortly thereafter, we tested P2X2/3 DKO mice in a similar protocol and observed strong preferences for both a fructose-paired CS+ flavor and for unflavored fructose; these data are presented below.

## 2. Materials and methods

### 2.1. Animals

Adult P2X2/3 DKO (6 male, 6 female) and WT mice (6 male, 6 female) were used. These mice were generated on a mixed C57BL/6 and 129Ola background [5;23]. As part of an unrelated experiment, the animals were food-restricted and given 1-min/day, 2-bottle tests with soybean oil emulsion (1.25 – 20% Intralipid) vs. water. The animals had no experience with sugar prior to the present experiment. The DKO and WT mice weighed 27.8 and 26.7 g, respectively, at the start of this experiment. The animals were singly housed in plastic tub cages with ad libitum access to chow (LabDiet 5001; PMI Nutrition International) and water in a room maintained at 22 °C with a 12:12 h light:dark cycle, except where noted. Experimental protocols were approved by the Institutional Animal Care and Use Committee at Brooklyn College and were performed in accordance with the National Institutes of Health Guidelines for the Care and Use of Laboratory Animals.

### 2.2. Test solutions

The solutions were prepared with food-grade fructose (Tate and Lyle, Honeyville Food Products, Rancho Cucamonga, CA), grape and cherry Kool Aid (Kraft Foods, Rye Brook, NY), and deionized water. The CS+ training solutions contained 8% fructose and 0.05% Kool-Aid and is referred to as CS+/F. The CS− solution was the other Kool-Aid flavor in water. For half the mice, the CS+ was cherry and the CS− was grape; for the remaining animals the CS flavors were reversed. In some two-bottle tests the CS+ flavor was presented in fructose (CS+/F) or in plain water (CS+). A 0.2% sodium saccharin solution (Sigma Aldrich, St. Louis, MO) was used in an initial screening test. In home cage training and testing the solutions were available through sipper spouts attached to 50-ml plastic tubes that were placed on top of the home cage. The sipper spouts were inserted through holes positioned 3.7 cm apart in a stainless-steel plate positioned to the right of the food bin, and the drinking tubes were fixed in place with clips. Fluid intakes were measured to the nearest 0.1 g by weighing the drinking bottles on an electronic balance interfaced to a laptop computer. Daily fluid spillage was estimated by recording the change in weight of two bottles that were placed on an empty cage, and intake measures were corrected by this amount.

Brief access two-bottle lick tests were conducted in plastic test cages as previously described [25]. Sipper spouts were attached to 50-ml glass tubes that were mounted on motorized bottle holders. The bottle holders positioned the spouts 1 mm in front of the cage at the start of a session and retracted them 1 min after the animal had emitted 10 licks. Licking behavior was monitored with electronic lickometers interfaced to a microcomputer.

### 2.3. Procedure

Prior to the start of the Intralipid experiment, the mice were given a 2-day screening test with 0.2% saccharin vs. water. At the beginning of the current experiment, the mice were water restricted overnight and given two 1-min, two-bottle lick tests, separated by 1 h, with 8% fructose vs. water (Test WR). They were then given ad libitum water and a restricted food ration (2 g) overnight followed the next day with two 1-min lick tests with 8% fructose vs. water (Test FR-1). The left-right positions of the sugar and water were alternated between the two tests conducted each day. Following 3 days of ad libitum food and water, the mice were then given a series of 24-h, two-bottle tests and one-bottle training sessions with flavored and unflavored fructose and water. In an initial 2-day pre-test (Test 0), CS+/F vs. CS− solutions were available. This was followed by four one-bottle training sessions with CS+/F and CS− presented on alternating days (CS+/F, CS−, CS+/F, CS−). Following a day of water only, the mice were given 2-day tests with CS+/F vs. CS− (Test 1), CS+ vs. CS− with both flavors in water only (Test 2), and then unflavored 8% fructose vs. unflavored water (Test 3). The left-right positions of the sugar and water bottles were alternated across the 2 days of each test. The mice were food restricted (2 g/day) and two days later were given two 1-min tests with fructose vs. water (Test FR-2) followed the next day with two 1-min tests with CS+/fructose vs. water (Test FR-3).

### 2.4. Data analysis

Fluid intakes in the 24-h tests were averaged over the 2 days of each two-bottle test or one-bottle training sessions with each fluid. Solution preferences in the two-bottle tests were expressed as percent intakes (e.g., fructose intake/total intake × 100). KO and WT group differences in solution intakes were evaluated using separate mixed model ANOVAs with group (genotype) and solution type as between-group and within-group factors, respectively. Significant interaction effects were evaluated using simple mean effects tests. Differences in solution preferences were evaluated using t-tests. One-min lick data were averaged over the two sessions of each test. Licks for each solution and percent preferences were evaluated as described above.

## 3. Results

In the initial saccharin screening test, the DKO mice consumed similar amounts of saccharin and water (3.6 vs. 3.8 g/day) whereas the WT mice consumed significantly more saccharin than water (7.8 vs. 1.6 g/day, Group × Solution interaction, F(1,22) = 15.0, p < 0.001). Consequently, the WT percent saccharin preference exceeded that of the DKO mice (79% vs. 50%, t(22) = 3.0, p < 0.001). When water restricted and given a 1-min choice test (Test WR) with 8% fructose vs. water, the DKO and WT mice did not differ in their total licks and did not differ in showing only a weak fructose preference, (59% vs. 60%) (Fig. 1). However, when food-restricted (Test FR-1), the WT mice licked more than DKO mice (F(1,22) = 67.1, p < 0.001) and, unlike DKO mice, licked more for fructose than water (Group × Solution interaction, F(1,22) = 67.1, p < 0.001). The fructose preference of the WT exceeded that of the DKO mice (93% vs. 60%, t(22) = 5.8, p < 0.001) (Fig. 1).

**Fig. 1.**
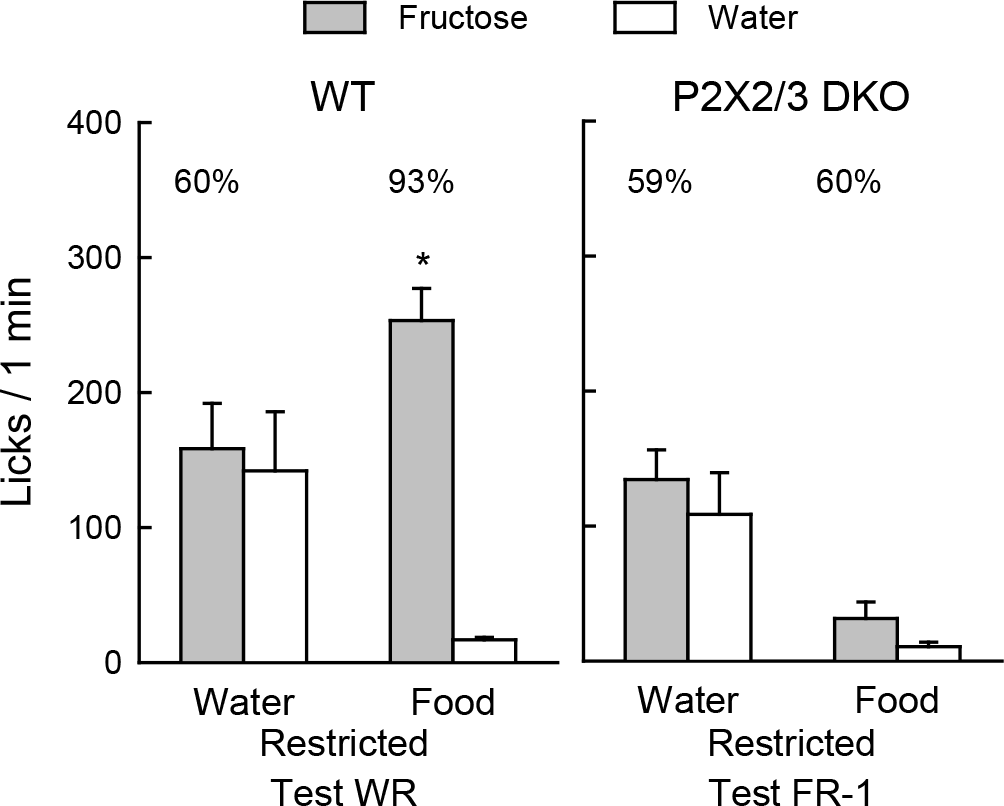
Mean (+SE) 8% fructose and water licks during 1-min two-bottle tests conducted under water restriction (Test WR) or food restriction (Test FR-1) for WT and P2X2/3 DKO mice. The mice had no prior experience with sugars. Numbers atop bars represent the mean percent preference for that sugar. Significant (p < 0.05) within group differences between solution licks indicated by *.

The results of the 24-h solution tests are presented in Fig. 2. In the two-bottle pretest (Test 0), both groups consumed more CS+/F than CS− (F(1,22) = 20.4, p < 0.001), although the WT mice tended to prefer the CS+/F more than the DKO mice (82% vs. 68%). This occurred because on the first day of Test 0 the DKO mice failed to prefer the CS+/F to CS− whereas the WT mice showed a strong preference (59% vs. 83%, t(22) = 2.2, p < 0.05); see Supplementary Fig. S1. On day 2, however, both DKO and WT mice displayed significant CS+/F preferences (74% vs. 80%) (Fig. S1). During the one-bottle training days the DKO mice consumed substantially more CS+/F than CS− whereas the WT mice consumed only slightly more, but the Group × Solution interaction was not significant. Following training, both groups consumed more CS+/F than CS− in Test 1 (F(1,22) = 30.7, p < 0.001). The DKO mice tended to consume more CS+ than WT and displayed a somewhat stronger CS+/F preference (88% vs. 80%), but these differences were not significant. In Test 2 both groups consumed more CS+ than CS− with both flavors presented in plain water (F(1,22) = 85.7, p < 0.001); the DKO and WT CS+ preferences were 93% and 82%, respectively, which did not significantly differ. In Test 3, the DKO and WT mice consumed more unflavored fructose than water (F(1,22) = 44.9, p < 0.001) and their sugar intakes and percent preferences (89% vs. 88%) did not differ.

**Fig. 2.**
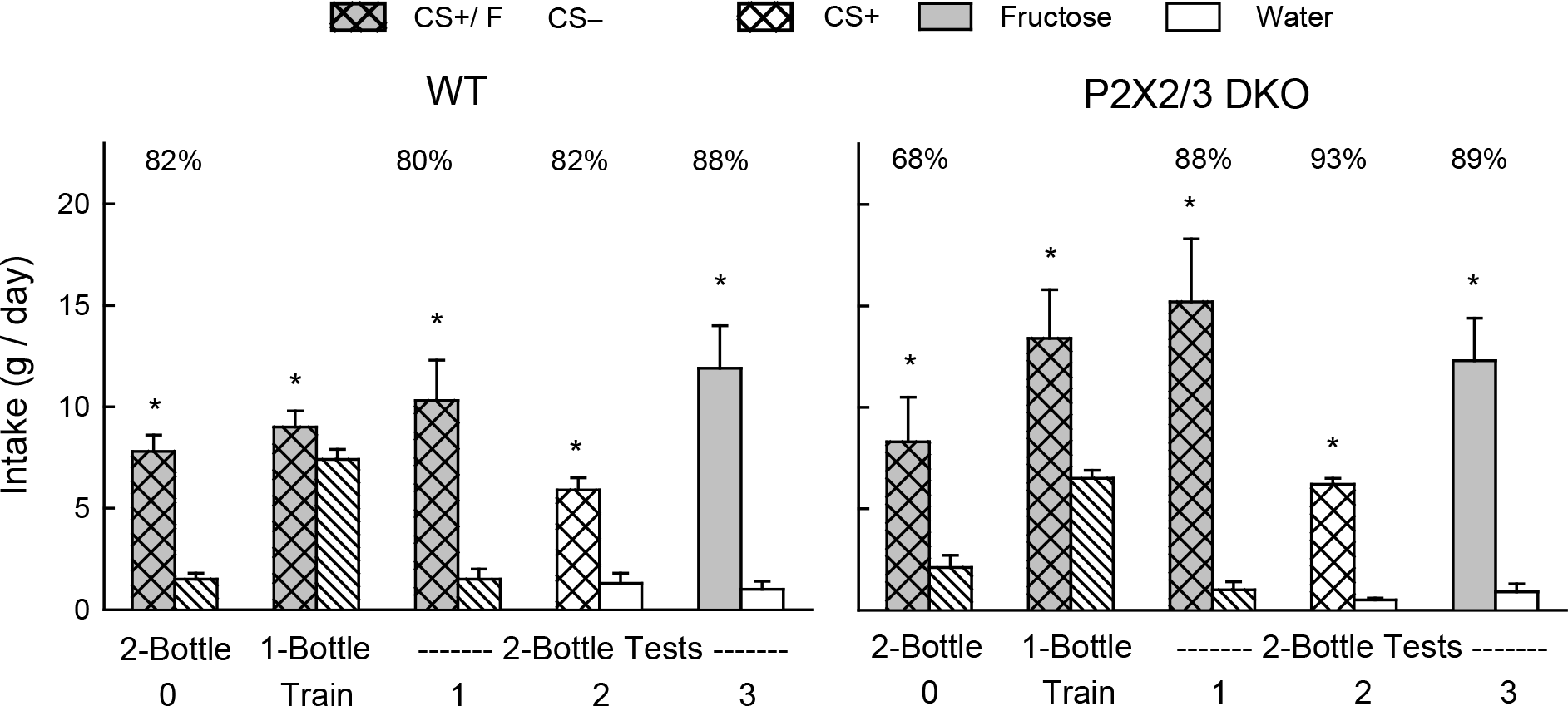
Mean (+SE) 24-h intake of CS+ fructose (CS+/F) and CS− water (CS−) during two-bottle Test 0 and Test 1 and one-bottle training sessions for WT and P2X2/3 DKO mice. In two-bottle Test 2 the mice were given CS+ flavored vs. CS− flavored water and in Test 3 unflavored 8% fructose vs. water. Numbers atop bars represent the mean percent preference for that solution. Significant (p < 0.05) within group differences between solution intakes indicated by *.

In the 1-min test with fructose vs. water (Test FR-2), which followed the 24-h test, the DKO mice were indifferent to fructose whereas the WT mice licked much more for fructose than water as they did in the initial 1-min test (Group × Solution interaction, F(1,22) = 37.2, p < 0.001). Consequently, the DKO mice showed a weaker fructose preference than WT mice (52% vs. 91% preference, t(22) =3.8, p < 0.001). When next given the choice of CS+ flavored fructose (CS+/F) and water (Test FR-3), the DKO mice licked more for CS+F than water, as did the WT mice, although they licked less for the CS+/F than did the WT mice (Group × Solution interaction, F(1,22) = 5.3, p < 0.05). The percent CS+/F preferences of the DKO and WT mice did not significantly differ at 83% and 95%, respectively.

**Fig. 3.**
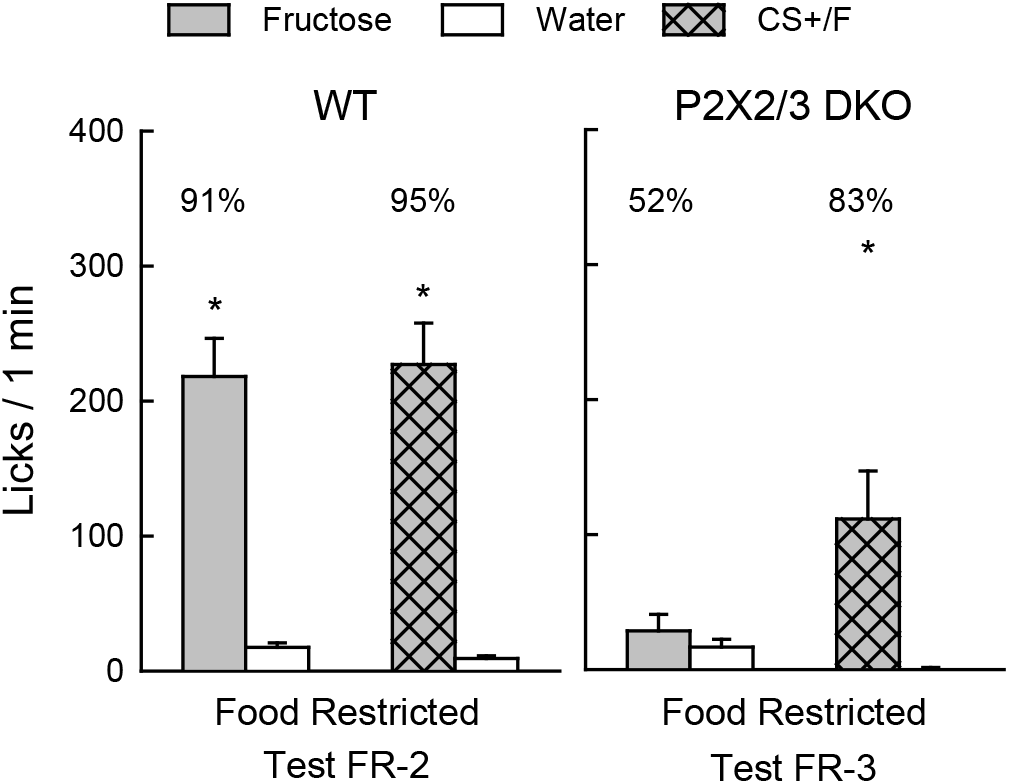
Mean (+SE) 8% fructose vs. water licks in 1-min two-bottle Test FR-2 and CS+ flavored 8% fructose vs. water in Test FR-3 conducted under food restriction for WT and P2X2/3 DKO mice. Numbers atop bars represent the mean percent preference for that sugar. Significant (p < 0.05) within group differences between solution licks indicated by *.

## 4. Discussion

The present results confirm the sweet taste ageusia of P2X2/3 DKO mice observed in prior studies [5;7;13]. The DKO mice, unlike WT mice failed to prefer 0.2% saccharin in 24-h tests or 8% fructose in 1-min sweetener vs. water tests (Test FR-1). Yet, in almost all 24-h tests with flavored or unflavored fructose the DKO mice displayed sugar preferences comparable to WT mice. The one exception was that the DKO mice failed to prefer fructose during the very first 24-h test but by the second day their sugar preference was comparable to WT mice. This finding, along with the initial 1-min fructose test results, demonstrates that the DKO mice were not inherently attracted to the sweet taste of the sugar but rather acquired a learned preference for the residual orosensory properties (i.e., odor, mouth feel) of the solution. The importance of sugar odor cues is indicated by results obtained with sweet ageusic T1R3 KO mice [27]. In particular, T1R2 KO mice were initially indifferent to 8% sucrose in a 24-h test, but after experience with more concentrated sugar solutions they subsequently displayed a near-total (95%) preference for 8% sucrose in a second test. Other T1R3 KO mice that were rendered anosmic by olfactory bulbectomy prior to the second sugar test displayed only a weak (67%) preference for 8% sucrose [27].

While experience-induced glucose and sucrose preferences have been observed in various types of ageusic KO mice (T1R3 KO, TRPM5 KO, Calhm1 KO, P2X2/3 DKO) [7;14;18;26], to date only P2X2/3 DKO mice show evidence of a fructose-experienced induced preference. In particular, whereas T1R3 KO and TRPM5 KO mice develop strong preferences for glucose or sucrose after 24-h experience with these sugars, they failed to prefer 8% fructose over water after experience with the sugar and they actually preferred water to 16% and 32% fructose [14;26]. TRPM5 KO mice also acquired a strong preference (96%) for a CS+ flavor mixed in 8% glucose but only a weak (64%) preference for a CS+ flavor mixed in 8% fructose [26]. These findings are consistent with the failure of B6 WT mice to learn to prefer a flavor paired with IG fructose infusions [12;24]. In contrast, the P2X2/3 DKO mice in the present study displayed a strong preference for the CS+ flavor alone as well as for the flavored (CS+/F) and unflavored fructose solutions in the 24-h tests. While P2X2/3 DKO and TRPM5 KO mice differ in their ability to acquire a fructose preference, they are similar in developing a strong preference for glucose and also preferring 8% glucose to 8% fructose in 24-h tests [14;22] (see Supplementary Fig. S2). P2X2/3 DKO and TRPM5 KO mice are also similar in acquiring preferences (79-85%) for a CS+ flavor mixed with monosodium glutamate (MSG) solutions, which is attributed to the postoral actions of the MSG [1;23].

In contrast to their 89% preference for unflavored fructose in 24-h Test 3, the P2X2/3 DKO mice failed to prefer fructose in the subsequent 1-min test (Test FR-2), just as they did in the initial 1-min test (Test FR-1) conducted while food deprived. It might be that their learned fructose preference was context specific, i.e., present in the home cage but not in the lick test cage. However, their preference for the CS+/F solution was not context specific but was displayed in both home and lick test cages. Instead, the home cage may provide stronger orosensory cues because the sipper tube is inserted into the cage while the sipper tube is presented outside a drinking slot in the lick test cage. In addition, delayed postoral appetition signals are available to guide the fluid preference in the 24-h tests but not during the 1-min tests.

Andres-Hernando *et al.* [2] recently reported that P2X2/3 DKO and WT mice display similar preferences for 15% fructose, and the present data indicate that this is also the case for a more dilute 8% fructose solution. However, Andres-Hernando *et al.* [2] observed a fructose preference on the very first test day, whereas the fructose preference did not appear until the second test day in the present experiment. Conceivably, the 15% sugar solution used in their study conditioned a more rapid acquired sugar preference than the 8% concentration used here. Yet, in another experiment we reported that T1R3 KO and TRPM5 KO mice did not display a preference for 34% sucrose until the second sugar vs. water test day, whereas the DKO mice in the Andres-Hernando *et al.* [2] study displayed a 15% sucrose preference on the first test day. Andres-Hernando *et al.* [2] hypothesized that P2X2/3 DKO mice show more rapid (i.e., day 1) sugar conditioning because, unlike T1R3 KO and TRPM5 KO mice, they are not missing the T1R3 and TRPM5 sugar sensing elements in the gut, which may contribute to postoral sugar appetition. However, this seems unlikely, particularly in the case of fructose, given that B6 WT mice with intact gut T1R3 and TRPM5 sugar sensing elements fail to acquire preferences for flavors paired with IG fructose infusions [12;24]. The first-day fructose preference in P2X2/3 DKO mice reported by Andres-Hernando *et al.* [2] is open to question because DKO mice tested with saccharin displayed a day 1 sweetener preference comparable to that of DKO mice tested with fructose. On subsequent days, the fructose preferences of the DKO mice increased while the saccharin preferences decreased.

Instead of differences in gut T1R3/TRPM5 signaling, the different fructose preferences displayed by TRPM5 KO and T1R3 KO mice vs. P2X2/3 DKO mice may be related to differences in their background genotypes. The TRPM5 KO and T1R3 KO background genotype is B6, which does not support postoral fructose appetition, while P2X2/3 DKO mice have a B6 × 129 hybrid genetic background. Conceivably, the B6 × 129 hybrid genotype enables fructose to have a postoral appetition action as observed in other strains (FVB, SWR) [8;20]. However, a recent study found that 129 mice and first generation B6 × 129 hybrid mice (B6:129:F1) showed no signs of fructose appetition [9]. It may be that B6:129:F1 mice do not exactly match the genetic background of P2X2/3 DKO and WT mice, which were subject to repeated inbreeding and cross-breeding (see [9]).

The major finding of the Andres-Hernando *et al.* [2] study was that P2X2/3 DKO mice, while displaying strong preferences for 15% fructose and other sugars, consumed less sugar than WT mice over a 30-week test period. Nevertheless, they consumed more chow and total calories and gained more weight than WT mice. The authors concluded that, independent of sweet taste, sugars can induce obesity and other aspects of the metabolic syndrome in mice. Somewhat different results were obtained with T1R3 KO and TRPM5 KO mice offered a 34% sucrose solution in addition to chow over a 38-day period [6]. In this case, the sugar-fed KO mice did not consume more total calories than WT mice, instead consuming the same or fewer calories. Also, the sugar-fed KO mice gained less body weight or body fat than WT mice. Furthermore, taste palatability influenced the carbohydrate intake and weight gain of T1R3 KO mice, because T1R3 KO mice offered a 34% Polycose (maltodextrin) solution which, unlike sucrose, is inherently attractive to the KO mice, overconsumed the Polycose and gained as much excess weight as did Polycose-fed WT mice. The findings of the different KO studies are difficult to compare given the different sugar concentrations (15% vs. 34%) and most importantly the study periods employed (30 weeks vs. 38 days). Future research should determine if long-term (∼30 weeks) access to sugar solutions induces adiposity and other metabolic changes in T1R3 KO and TRPM5 KO mice similar to those of P2X2/3 DKO mice.

## Acknowledgement

This research was supported by grant DK-31135 from the National Institute of Diabetes and Digestive and Kidney Diseases. The P2X2/3 DKO and WT mice were supplied by the Rocky Mountain Taste & Smell Center that was supported by the National Institute on Deafness and Other Communication Disorders Grant P30 DC004657.

## Supplementary Figures

**Fig. S1.**
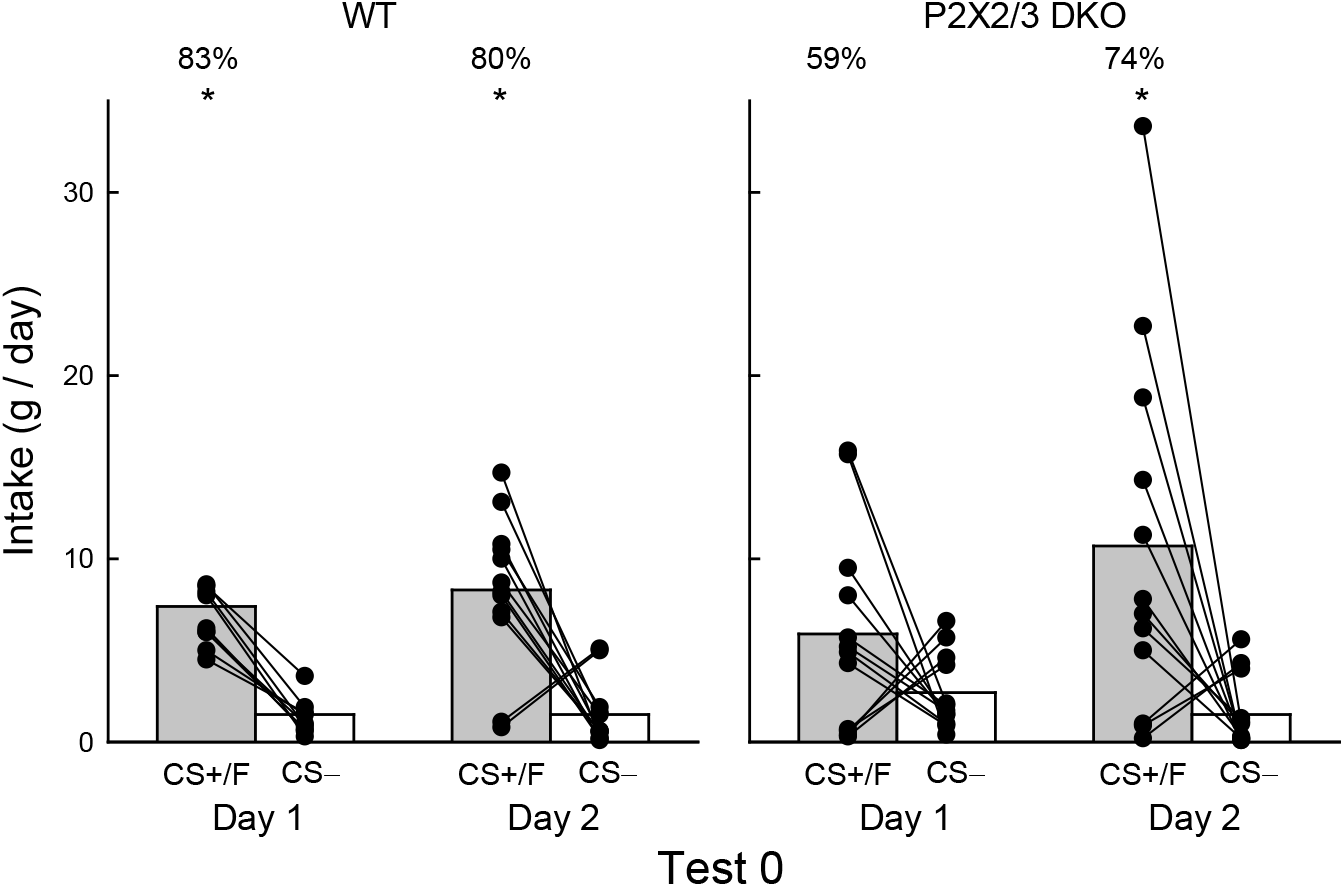
Test 0 Daily Data. Group mean (bars) and individual subject (lines) 24-h intakes of CS+ fructose (CS+/F) and CS− water (CS−) during days 1 and 2 of two-bottle Test 0 for WT and P2X2/3 DKO mice. Numbers atop bars represent the mean percent preference for CS+/F. Significant (p < 0.05) within group differences between solution intakes indicated by *.

**Fig. S2.**
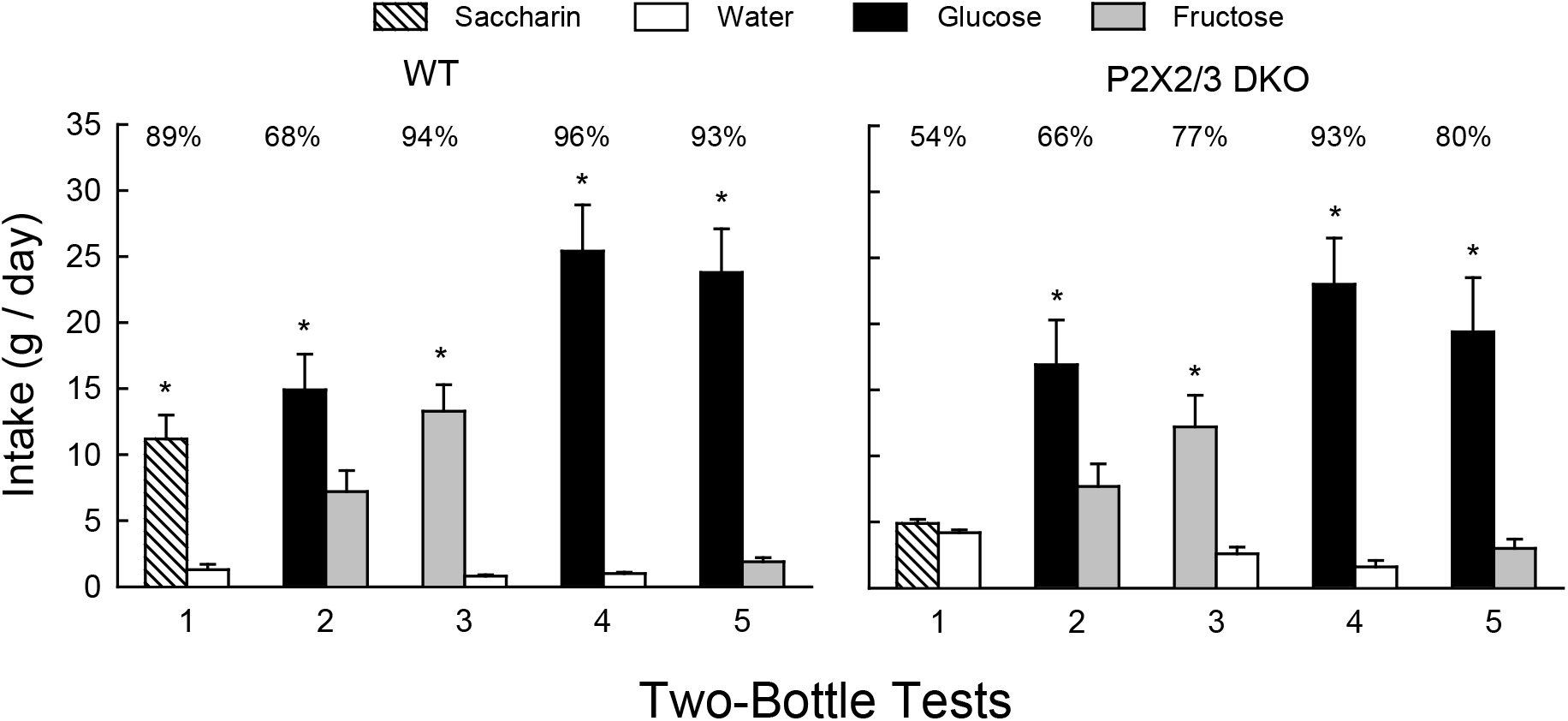
P2X2/3 DKO Fructose and Glucose Preference Data. Mean (+SE) 24-h intakes during 2-day tests with 0.2% saccharin vs. water (Test 1), 8% glucose vs. 8% fructose (Test 2), 8% fructose vs. water (Test 3), 8% glucose vs. water (Test 4) and 8% glucose vs. 8% fructose (Test 5) for WT (n = 11) and P2X2/3 DKO mice (n=11). Numbers atop bars represent the mean percent preference for that solution. Significant (p < 0.05) within group differences between solution intakes indicated by *. The mice had prior 24-h experience with saccharin, Polycose [13], sucrose and fructose prior to Test 1, but no prior experience with glucose. The preferences of the P2X2/3 DKO mice for glucose over water (Test 4, 93%) and glucose over fructose (Test 5, 80%) are similar to the preferences (85%) displayed by TRPM5 KO mice [14;22]. In contrast, whereas the P2X2/3 DKO preferred fructose to water by 77% (Test 3), TRPM5 KO mice displayed only a 55% preference [14].

## Notes

### Competing Interest Statement

The authors have declared no competing interest.

### Summary of Updates

Figure 2 revised

